# Molecular basis of immune evasion by the delta and kappa SARS-CoV-2 variants

**DOI:** 10.1101/2021.08.11.455956

**Authors:** Matthew McCallum, Alexandra C. Walls, Kaitlin R. Sprouse, John E. Bowen, Laura Rosen, Ha V. Dang, Anna deMarco, Nicholas Franko, Sasha W Tilles, Jennifer Logue, Marcos C. Miranda, Margaret Ahlrichs, Lauren Carter, Gyorgy Snell, Matteo Samuele Pizzuto, Helen Y. Chu, Wesley C. Van Voorhis, Davide Corti, David Veesler

## Abstract

Worldwide SARS-CoV-2 transmission leads to the recurrent emergence of variants, such as the recently described B.1.617.1 (kappa), B.1.617.2 (delta) and B.1.617.2+ (delta+). The B.1.617.2 (delta) variant of concern is causing a new wave of infections in many countries, mostly affecting unvaccinated individuals, and has become globally dominant. We show that these variants dampen the in vitro potency of vaccine-elicited serum neutralizing antibodies and provide a structural framework for describing the impact of individual mutations on immune evasion. Mutations in the B.1.617.1 (kappa) and B.1.617.2 (delta) spike glycoproteins abrogate recognition by several monoclonal antibodies via alteration of key antigenic sites, including an unexpected remodeling of the B.1.617.2 (delta) N-terminal domain. The binding affinity of the B.1.617.1 (kappa) and B.1.617.2 (delta) receptor-binding domain for ACE2 is comparable to the ancestral virus whereas B.1.617.2+ (delta+) exhibits markedly reduced affinity. We describe a previously uncharacterized class of N-terminal domain-directed human neutralizing monoclonal antibodies cross-reacting with several variants of concern, revealing a possible target for vaccine development.

---

The ongoing spread of SARS-CoV-2, the causative agent of the COVID-19 pandemic, results in the continued emergence of variants. The B.1.351 (beta, β) variant of concern was originally described in South Africa and remains the isolate associated with the greatest magnitude of immune evasion, as measured by reduced neutralizing antibody (Ab) titers in vitro (*1–3*). Conversely, the B.1.1.7 (alpha, α) variant of concern, which was first detected in the United Kingdom, has a modest impact on neutralizing Ab titers but a marked enhancement in ACE2 receptor binding affinity and transmissibility which led to worldwide dominance in the early months of 2021 (*2, 4*).

The SARS-CoV-2 spike (S) glycoprotein is exposed at the surface of the virus and mediates entry into host cells. S is the main target of neutralizing antibodies and the focus of most vaccines (*5, 6*). The S glycoprotein is subdivided into two functional subunits, designated S_1_ and S_2_, that interact non-covalently after proteolytic cleavage by furin during synthesis (*5, 7, 8*). The S_1_ subunit contains the receptor-binding domain (RBD), which engages the receptor ACE2 (*5, 7, 9, 10*), and the N-terminal domain (NTD) that recognizes attachment factors (*11–13*). The S_2_ subunit contains the fusion machinery and undergoes large-scale conformational changes to drive fusion of the virus and host membranes (*14*), enabling genome delivery and initiation of infection. Abs that bind to specific sites on the RBD (*15–22*), the NTD (*23–26*), or the fusion machinery stem helix (*27*– *31*) interfere with receptor attachment or membrane fusion. Serum neutralizing Ab titers are a correlate of protection against SARS-CoV-2 in non-human primates (*32–35*).

In late 2020, B.1.617 variants including B.1.617.1 (kappa, κ) and B.1.617.2 (delta, δ) were first detected in India and caused devastating epidemics before spreading globally (*36, 37*). The variant S harbors T95I, G142D, E154K, L452R, E484Q, D614G, P681R and Q1071H substitutions whereas the B.1.617.2 variant S carries T19R, G142D, E156G, L452R, T478K, and D950N substitutions and a deletion of residues 157 and 158 (157-158del) (**Table S1**). Most of these mutations localize to the RBD and NTD which are the major targets of neutralizing Abs in convalescent and vaccinated individuals, raising concerns about the efficacy of available vaccines and therapeutic monoclonal Abs (mAbs) against these variants. Moreover, the K417N mutation was detected in the B.1.617.2 lineage, known as the B.1.617.2+ (delta plus, δ+) variant, which is shared with the B.1.351 (beta, β) variant of concern and was previously shown to reduce neutralization potency of some monoclonal Abs (*2, 38*).

To evaluate the effect on neutralization of the mutations in the B.1.617.1, B.1.617.2, and B.1.617.2+ S glycoproteins, we compared vaccine-elicited serum neutralizing activity against ancestral G614 S and these three variant S pseudoviruses. We employed a vesicular stomatitis virus (VSV) pseudotyping system (*39*) with ACE2-expressing HEK293T as target cells (*40*). Plasma was used from 13, 15, and 8 individuals who received two doses of Pfizer/BioNtech BNT162b2, two doses of Moderna mRNA-1273, or a single dose of Janssen Ad26.COV2.S, respectively (**Table S2**). Geometric mean neutralizing Ab titers (GMTs) for the Pfizer/BioNtech BNT162b2-elicited plasma were respectively reduced 3-, 3-, and 5-fold for B.1.617.1, B.1.617.2, and B.1.617.2+ S (GMTs 84, 110, and 62), compared to G614 S (GMT: 280) (**Fig. 1A, Fig S1-S2**). Moderna mRNA-1273-elicited plasma GMTs were reduced 3-, 2-, and 9-fold for B.1.617.1,, and B.1.617.2+ S (GMTs 270, 410, and 94), respectively, compared to G614 S (GMT: 850) (**Fig. 1B, Fig S1-S2**). The average neutralization potency of the Janssen Ad26.COV2.S-elicited plasma was reduced 5-, 4-, and ≥6-fold for B.1.617.1, B.1.617.2, and B.1.617.2+ S (GMTs 9.8, 12, and 8.4), respectively, compared to D614G S (GMT: 47) (**Fig. 1C, Fig S1-S2**). These data demonstrate that all three variants lead to reductions in neutralization potency from vaccine-elicited Abs with B.1.617.2+ causing the greatest decrease, on par with what was observed for the B.1.351 (beta) variant of concern (*1, 3*). Furthermore, polyclonal Abs from half of the Janssen-vaccinated individuals evaluated completely lost their ability to neutralize one or multiple variants in our assay, likely as a result of the moderate GMTs against G614 S pseudovirus (*41*).

**Figure 1.**
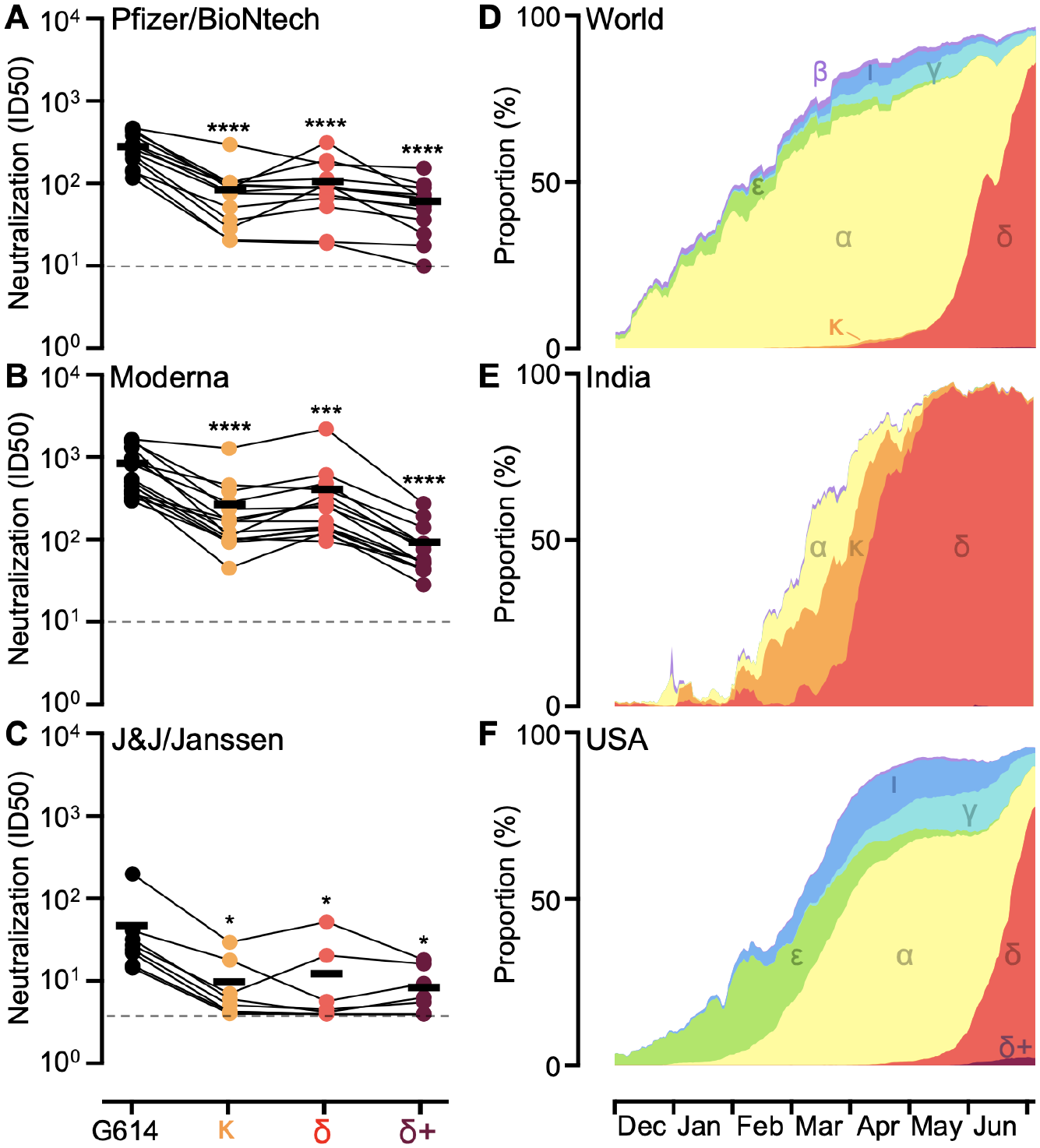
Neutralization of pseudotyped viruses harboring SARS-CoV-2 G614 S, B.1.617.1 S, B.1.617.2 S, or B.1.617.2+ S and incidence of variants. (A-C) Pairwise connected 50% inhibition concentrations (IC50s) for each individual against each variant for Pfizer/BioNtech BNT162b2 (A), Moderna mRNA-1273 (B), or J&J/Janssen Ad26.COV2.S (C) vaccinee sera (**Fig. S1** and **Table S1**). Data are an average of n = 2 replicates and are representative of at least two independent assays. Dashed line indicates the limit of detection for the assay. Means (shown as thick black horizontal lines) were compared against G614 by two-way ANOVA (Dunnet’s test); *, p<0.05; ***, p<0.001; ****, p<0.0001. (D-F) Incidence of variants of concern and variants of interest as a proportion of viruses sequenced in the world (D), India (E), and the USA (F) deposited to GISAID (analyzed using outbreak.info) from December 1, 2020 to July 4, 2021. B.1.1.7 (alpha, α), B.1.351 (beta, β), P.1 (gamma, γ), B.1.617.2 (delta, δ) including AY.3, B.1.526 (iota, ι), B.1.427/B.1.429 (epsilon, ε), B.1.617.1 (kappa, κ), and B.1.617.2+ (delta+, δ+) including AY.1 and AY.2 are shown in yellow, light purple, cyan, red, blue, green, orange, and dark purple, respectively.

**Figure S1:**
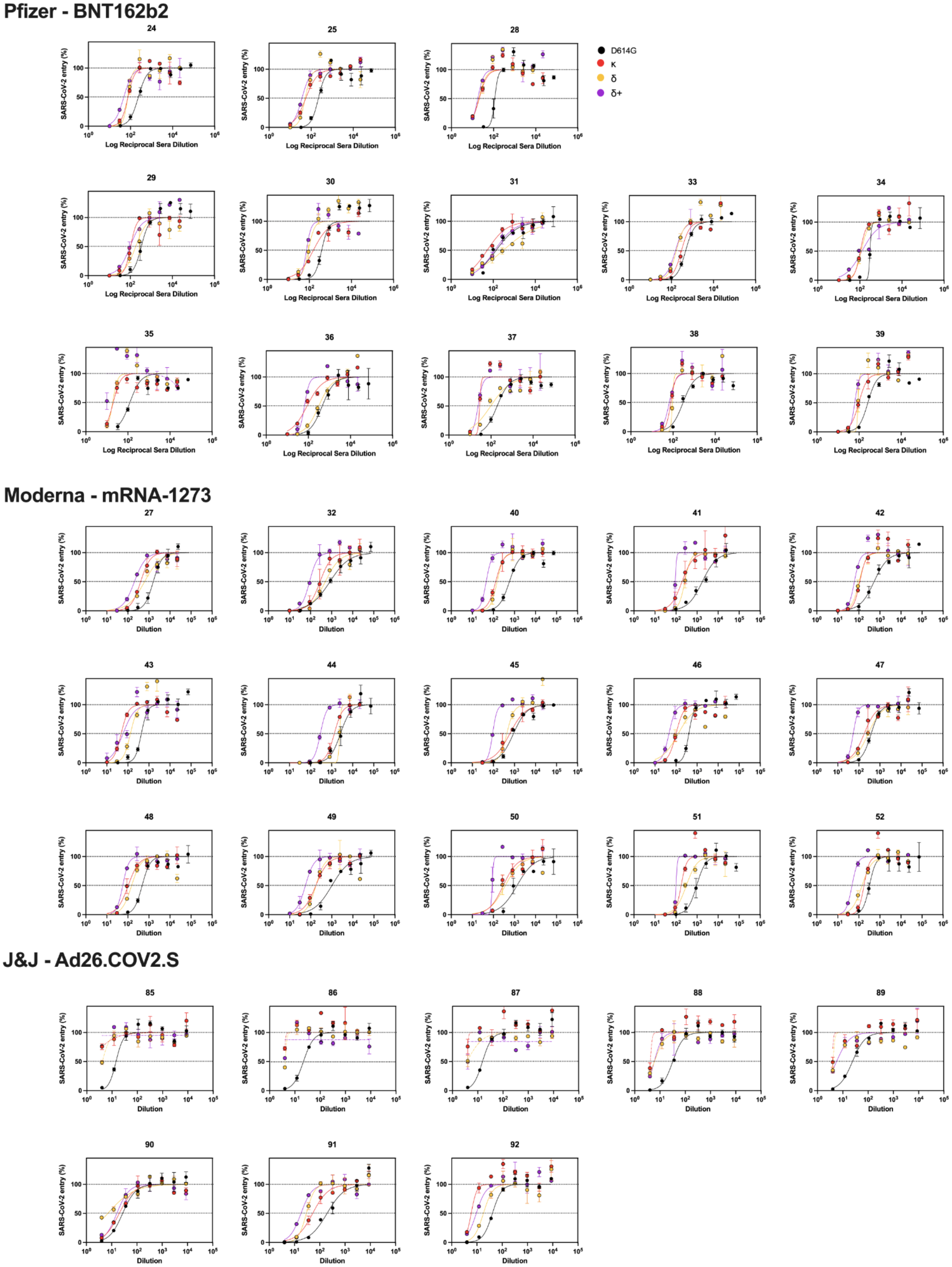
Neutralization of vaccinated human sera. Related to Fig. 1A-C. Normalized neutralization data points and fits of individual sera from Pfizer, Moderna, and J&J against D614G, κ, δ, and δ+ VSV pseudovirus.

**Figure S2:**
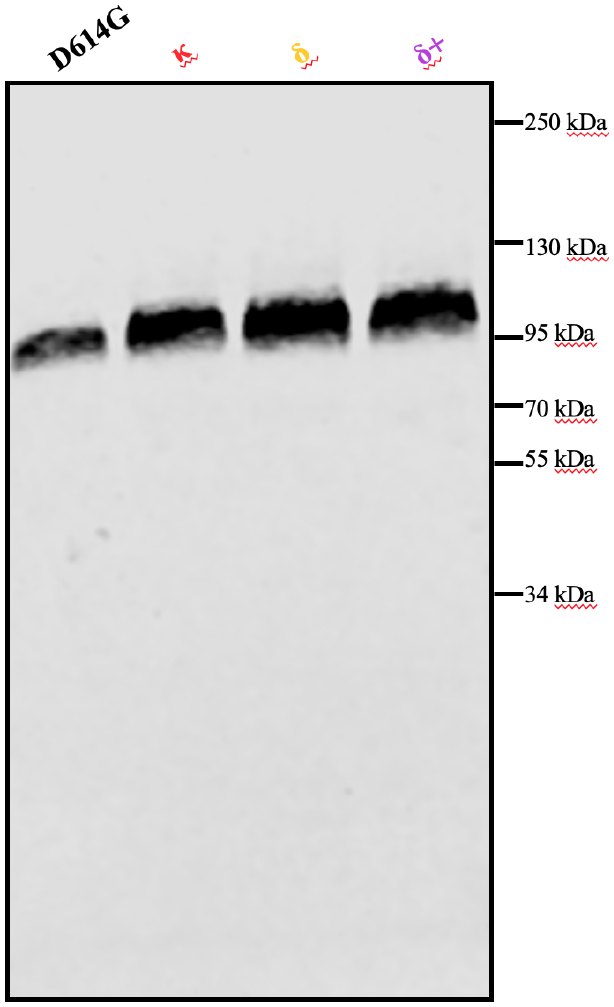
Western blot comparing S of pseudotypes used for neutralization. VSV pseudotype G614, κ, δ, and δ+ S visualized by western blotting showing relative amounts of S used in the neutralization assays.

Although the B.1.617.1 and B.1.617.2+ variants have greater ability to evade vaccine-elicited Ab neutralization than B.1.617.2 – a feature that might increase opportunities for breakthrough infections (*37, 42, 43*) – the B.1.617.2 variant became dominant worldwide by June 2021 (**Fig 1D-F**). B.1.617.1 and B.1.617.2 sequences were first observed in late 2020 whereas B.1.617.2+ sequences were first detected in April 2021. As of August 9, 2021, 6,017 B.1.617.1, 376,967 B.1.617.2, and 1,626 B.1.617.2+ sequenced genomes were deposited in GISAID (**Table S1**). The high incidence of B.1.617.2 is consistent with recent studies showing that the B.1.617.2 variant has enhanced transmissibility, replication kinetics and viral loads in oropharyngeal and nose-throat swabs of infected individuals relative to the ancestral virus and other variants (*37, 44*).

We determined electron cryomicroscopy (cryoEM) structures of the B.1.617.1 and B.1.617.2 S glycoproteins to gain insights into their reduced antibody sensitivity and provide a structural framework to understand the role of shared and distinct mutations. The B.1.617.1 S structure – harboring five of the HexaPro stabilizing mutations (*45*), a native furin cleavage site, and engineered in the closed conformation with S383C/D985C mutations (*46*) – was determined in complex with S309 (an RBD-targeted neutralizing mAb (*15*)) and S2L20 (an NTD-targeted non-neutralizing mAb (*23, 47*)) at 2.4 Å resolution (**Fig 2A, Fig S3A-C, Table S3**). Local classification and refinements of S309 bound to the B.1.617.1 RBD were used to account for conformational dynamics and improve local resolution of this region to 3.3 Å (**Fig 2A,C and Fig S3A**). The B.1.617.2 S structure – harboring some of the HexaPro (*45*) and VFLIP (*48*) stabilizing mutations and a native furin cleavage site – was obtained in complex with S2M11 (an RBD-targeted neutralizing mAb (*19*)) and S2L20 at 2.4 Å resolution (**Fig 2B, Fig S3D-F, Table S3**).

**Figure 2.**
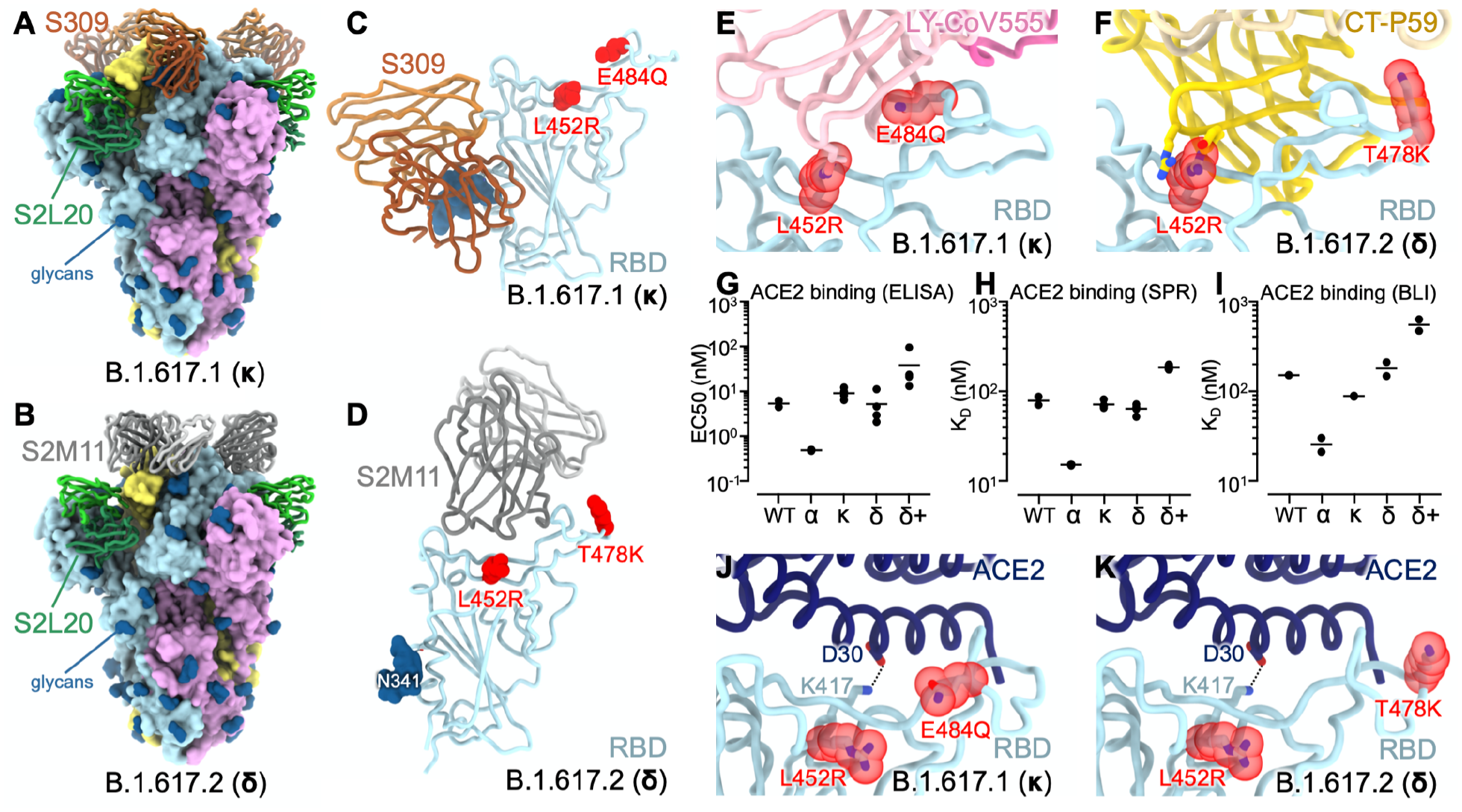
CryoEM structures of the SARS-CoV-2 B.1.617.1 and B.1.617.2 S ectodomain trimers and analysis of ACE2 binding. **(A-B)** Structure of the B.1.617.1 (A) and B.1.617.2 (B) S trimer (surface rendering) bound to the S2L20 and S309 (A) or S2M11 (B) Fabs (ribbons). SARS-CoV-2 S protomers are colored pink, cyan, and gold, whereas the S2L20 Fab heavy and light chains are colored dark and light green, respectively. The S309 Fab heavy and light chains are colored dark and light orange, respectively (A). The S2M11 Fab heavy and light chains are colored dark and light gray, respectively (B). Only the Fab variable domains are resolved and therefore modeled in the map. N-linked glycans are rendered as dark blue spheres. (**C**) Zoomed in view of the S309-bound B.1.617.1 RBD with L452R and E484Q shown as red spheres. (**D**) Zoomed in view of the S2M11-bound B.1.617.2 RBD with L452R and T478K shown as red spheres. (**E**) Superimposition of the LY-CoV555–bound SARS-CoV-2 RBD structure (PDB 7KMG) on the SARS-CoV-2 B.1.617.1 S cryoEM structure show that L452R would clash with the mAb and E484Q would disrupt electrostatic interactions. (**F**) Superimposition of the CT-P59–bound SARS-CoV-2 RBD structure (PDB 7CM4) on the SARS-CoV-2 B.1.617.2 S cryoEM structure show that L452R would sterically clash with the mAb. (**G**) Enzyme-linked immunosorbant assay (ELISA) binding analysis of the SARS-CoV-2 wildtype, B.1.1.7 (α), B.1.617.1 (κ), B.1.617.2 (δ), and B.1.617.2+ (δ+) RBDs to immobilized human ACE2 ectodomain (residues 1-615) shown as 50% effective concentrations (EC_50_). Data from two biological replicates are shown with 2-4 technical replicates each. (**H**) Surface plasmon resonance (SPR) binding affinity analysis of the human ACE2 ectodomain (residues 1-615) for immobilized biotinylated wildtype, B.1.1.7, B.1.617.1, B.1.617.2, and B.1.617.2+ RBDs. Data from two biological replicates are shown with 2-6 technical replicates each. (**I**) Biolayer Interferometry (BLI) binding analysis of the human ACE2 ectodomain (residues 1-615) to immobilized biotinylated SARS-CoV-2 wildtype, B.1.1.7, B.1.617.1, B.1.617.2, and B.1.617.2+ RBDs. Data from two biological replicates are shown with 1-2 technical replicates each. (**J-K**) Superimposition of the ACE2-bound SARS-CoV-2 RBD structure (PDB 6VW1) on the SARS-CoV-2 B.1.617.1 (J) and B.1.617.2 (K) S cryoEM structures show that L452R and T478K point away from the interface with ACE2, while K417 contacts D30 from ACE2.

**Fig. S3.**
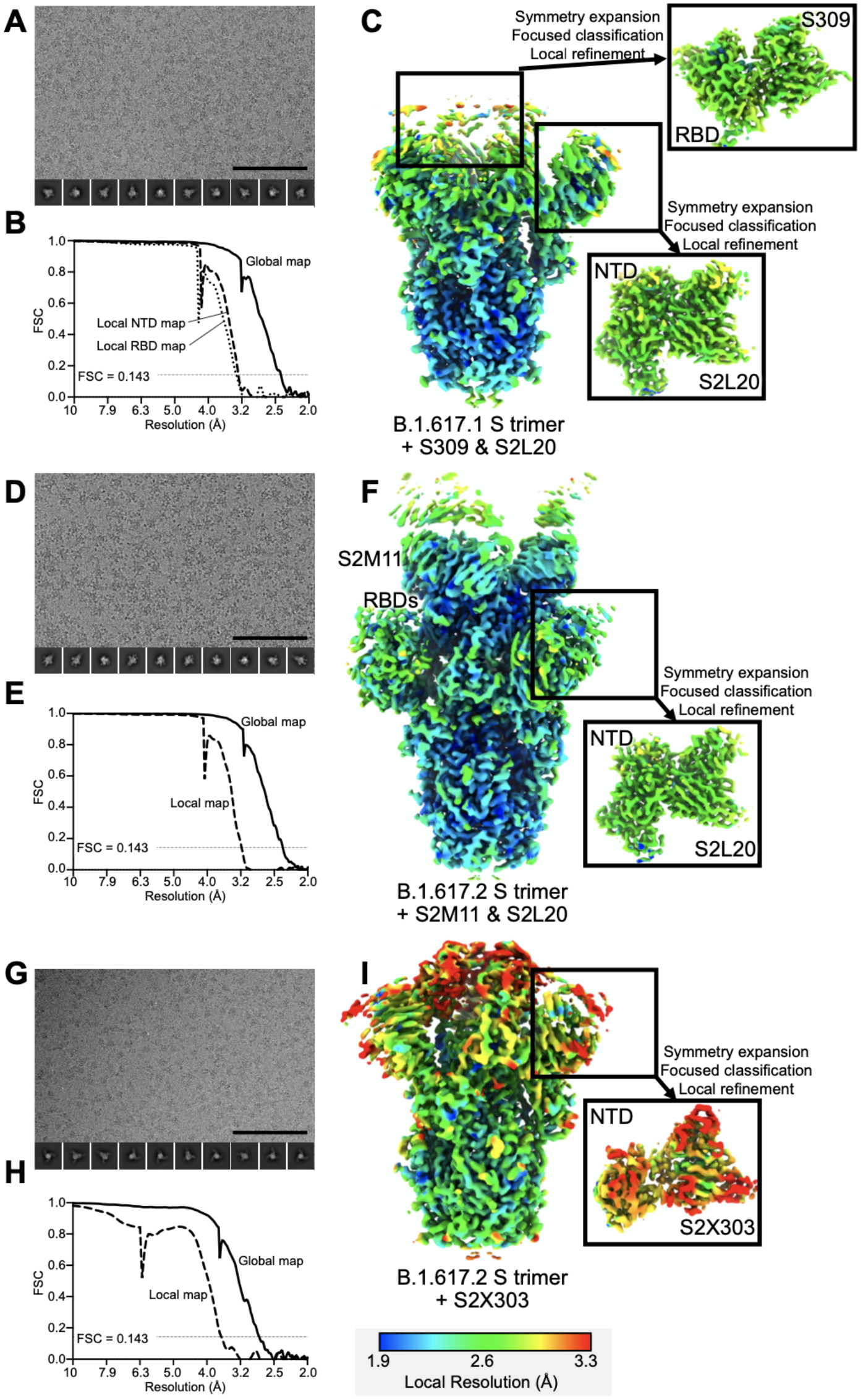
CryoEM data processing of SARS-CoV-2 B.1.617.1 S ectodomain trimer bound to S2L20 and S309 (A-C), B.1.617.2 S ectodomain trimer bound to S2L20 and S2M11 (D-F), and B.1.617.1 S ectodomain trimer bound to S2X303 (G-I). **(A, D, and G)** Representative electron micrograph (bottom right, scale bar: 100 nm) and 2D class averages (bottom) are shown for the indicated particles embedded in vitreous ice. **(B, E, and H)** Gold-standard Fourier shell correlation curves with the 0.143 cutoff indicated by a horizontal dashed line. **(C, F, and I)** Unsharpened maps colored by local resolution calculated using cryoSPARC for the whole reconstruction and the locally refined reconstructions of NTD- or RBD-bound Fab variable domains (insets).

The RBD is the main target of serum neutralizing activity in convalescent and vaccinated individuals and comprises several antigenic sites recognized by neutralizing Abs with a range of neutralization potencies and breadth (*15–18, 49–51*). The B.1.617.1 S and B.1.617.2 S structures resolve the complete RBD and provide high-resolution blueprints of the residue substitutions found in these two variants (**Fig 2C-D**). In both structures, the L452R side chain, which is part of antigenic site Ib (*18*), is oriented identically (and modeled in the same rotameric configuration) to what we previously observed in the B.1.427/B.1.429 S structure (*47*) (**Fig 2E-F**). The L452R mutation reduces neutralization mediated by some clinical mAbs, such as bamlanivimab (LY-CoV555) and regdanvimab (CT-P59), due to steric alteration of this antigenic site that is incompatible with binding (**Fig 2E-F**) (*47*). The B.1.617.1 E484Q RBD mutation is located within the receptor-binding motif, at the boundary between antigenic sites Ia/Ib (*18*), and could affect neutralization from mAbs recognizing both subsites. However, the E484Q mutation is conservative as it substitutes the side chain carboxylic group with an amide group through replacement of an oxygen with a nitrogen atom. Residue 484 is involved in the epitopes recognized by bamlanivimab (LY-CoV555) (*20, 52*) and casirivimab (REG10933) (*53, 54*), and both mAbs have dampened binding or neutralization potency against E484Q-harboring mutants likely due to disruption of electrostatic interactions (*52, 54*). The B.1.617.2 T478K RBD mutation is also located at the boundary between antigenic sites Ia/Ib but does not affect binding to mAbs that have received an emergency use authorization in the US (*52, 55, 56*) (**Fig 2D**).

E484Q, L452R, and T478K mutations are part of antigenic site I which we previously showed to be immunodominant (*16–18*). Both the B.1.351 (beta) and P.1 (gamma) variants of concern harbor the E484K mutation which is associated with a reduction of vaccine-elicited neutralizing Ab titers and mAb neutralization (*38, 41, 57*). The E484Q mutation has also been reported to reduce serum neutralizing Ab titers alone or in combination with the L452R mutation (*41, 54, 58*). The effect of the T478K mutation on polyclonal Ab responses has not been characterized as thoroughly as the L452R and E484Q substitutions. The recurrent emergence of mutants carrying residue substitutions at position 452 and 484 underscores the apparent hyperfocusing of neutralizing antibody responses at these antigenic subsites and provides a possible explanation for their acquisition in multiple SARS-CoV-2 lineages.

We set out to assess the impact of the RBD mutations present in the B.1.617.1, B.1.617.2, and B.1.617.2+ on binding affinity for the ACE2 receptor to assess potential impact on a component of viral transmissibility. The B.1.617.1 (L452R/E484Q) and B.1.617.2 (L452R/T478K) RBDs recognized immobilized ACE2 with roughly similar efficiency to the wildtype RBD, as observed by ELISA (**Fig 2G, Fig S4, Table S4**). ACE2 binding to the B.1.617.2+ (K417N/L452R/T478K) and B.1.1.7 (N501Y) RBDs was respectively much weaker and much tighter than for any other variants evaluated in our assay, in agreement with the enhanced affinity conferred by N501Y (*4, 59*) (**Fig 2G, Table S4**). We confirmed these results using surface plasmon resonance (**Fig 2H, Fig S4, Table S4**) and biolayer interferometry (**Fig 2I, Fig S4X, Table S4**) binding analysis of the monomeric human ACE2 ectodomain to immobilized RBDs, indicating that the B.1.617.1 and B.1.617.2 RBDs interact with ACE2 with roughly comparable affinity to the wildtype RBD whereas the B.1.617.2+ RBD is severely attenuated. These data concur with evaluation of the effect of individual residue mutations on ACE2 binding using deep-mutational scanning of yeast-displayed RBDs (*59*), the positioning of the R452 and K478 side chains away from the ACE2-binding interface, and the conservative nature of the E484Q substitution (**Fig 2J-K**). Finally, these data point to the key contribution of the salt bridge formed between the K417_SARS-CoV-2_ and D30_ACE2_ side chains for receptor engagement (**Fig 2J-K**), in agreement with previous studies (*60, 61*). Since none of the and B.1.617.2 RBD mutations increase ACE2 binding markedly and all of them reside in the most immunogenic antigenic site, we propose that they emerged mainly as a result of antibody-mediated selective pressure to reduce immune recognition.

**Figure S4:**
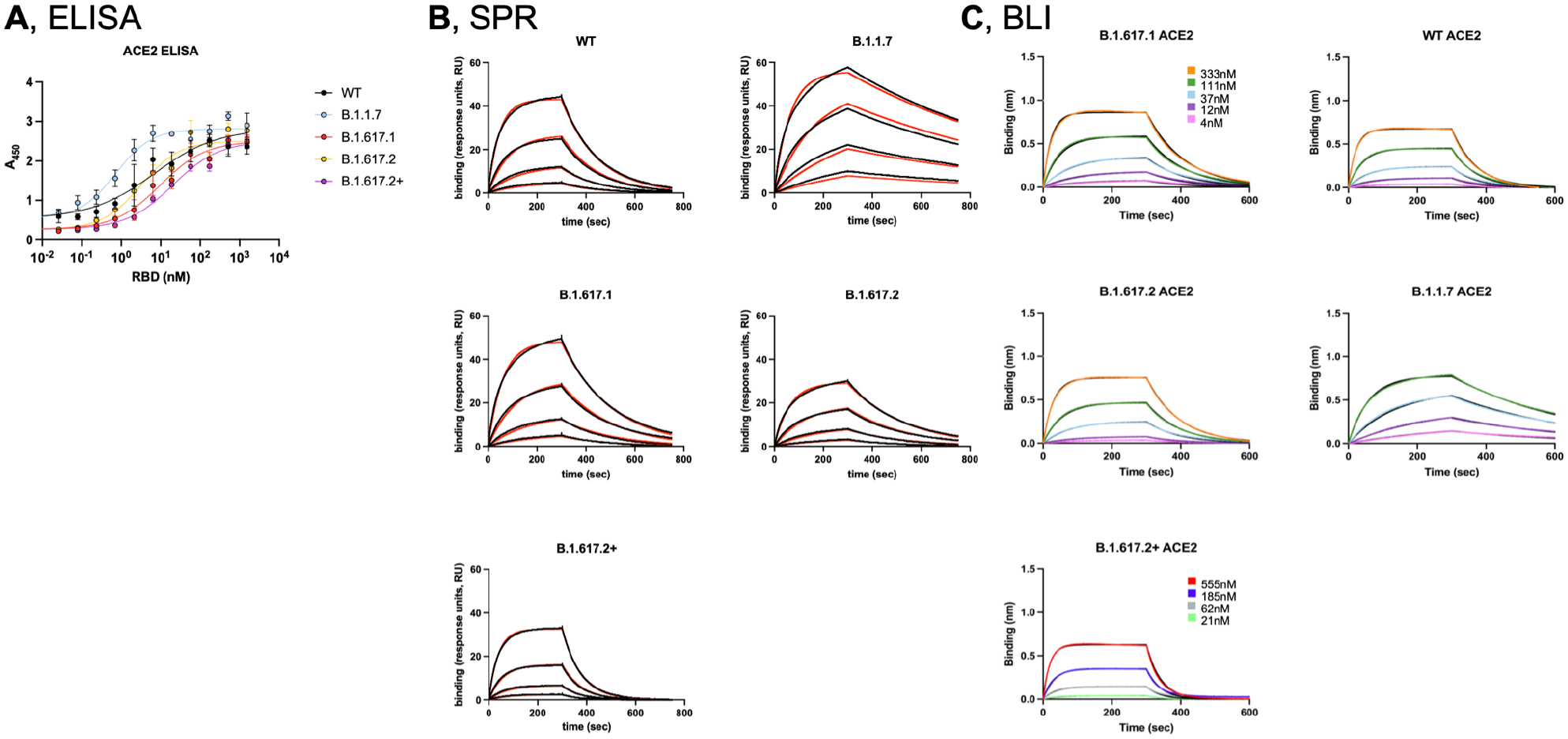
Binding kinetics of RBDs to ACE2 using ELISA, SPR, and BLI. (**A)** Binding of the SARS-CoV-2 wildtype, B.1.1.7, B.1.351, B.1.617.1, B.1.617.2, and B.1.617.2+ RBDs to immobilized human ACE2 ectodomain analyzed by ELISA. (**B**) Binding human ACE2 to the immobilized SARS-CoV-2 wildtype, B.1.1.7, B.1.617.1, B.1.617.2, and B.1.617.2+ RBDs analyzed by SPR. (**C**) Binding human ACE2 to the immobilized SARS-CoV-2 wildtype, B.1.1.7, B.1.617.1, B.1.617.2, and B.1.617.2+ RBDs analyzed by BLI. Binding kinetics shown in Table S3.

Although multiple antigenic sites are present at the surface of the NTD, a single supersite of vulnerability is known to be targeted by neutralizing Abs elicited upon infection and vaccination (*23–25*). This antigenic supersite (designated site i) comprises the NTD N-terminus (residues 14-20), a β-hairpin (residues 140-158), and a loop (residues 245-264). To improve the cryoEM map resolution of the B.1.617.1 and B.1.617.2 NTDs bound by S2L20 and visualize the structural changes associated with their respective constellation of mutations, we used focused 3D classification and local refinement. Two of the three B.1.617.1 NTD mutations, G142D and E154K, map to the supersite β-hairpin (**Fig 3A, Table S1)** and we have previously shown that G142D abrogates binding of 3 out of 5 NTD-specific neutralizing mAbs tested (*23*). The B.1.617.1 T95I substitution occurs outside the antigenic supersite and is unlikely to contribute to immune evasion significantly (**Fig 3A)**. All of the B.1.617.2 mutations are found within the antigenic supersite: T19R, G142D, E156G, and 157-158del (**Fig 3B)**. The T19R substitution abrogates the glycosylation sequon at position N17, as supported by the lack of a resolved glycan at this position in the cryoEM map. We previously showed that T19A, which also removes the N17 glycosylation sequon, decreased binding to 4 out 5 NTD neutralizing mAbs tested (*23*). Residues 156-158 participate in the supersite β-hairpin and their mutation/deletion in the B.1.617.2 NTD lead to striking structural remodeling: residues 151-159 adopt an alpha-helical conformation whereas this segment is β-stranded in the absence of this mutation/deletion (**Fig 3B)**. Based on these findings, we hypothesized that B.1.617.1 and B.1.617.2 variant S glycoproteins would escape recognition by most neutralizing NTD Abs (**Fig 3C & 3D)**.

**Figure 3.**
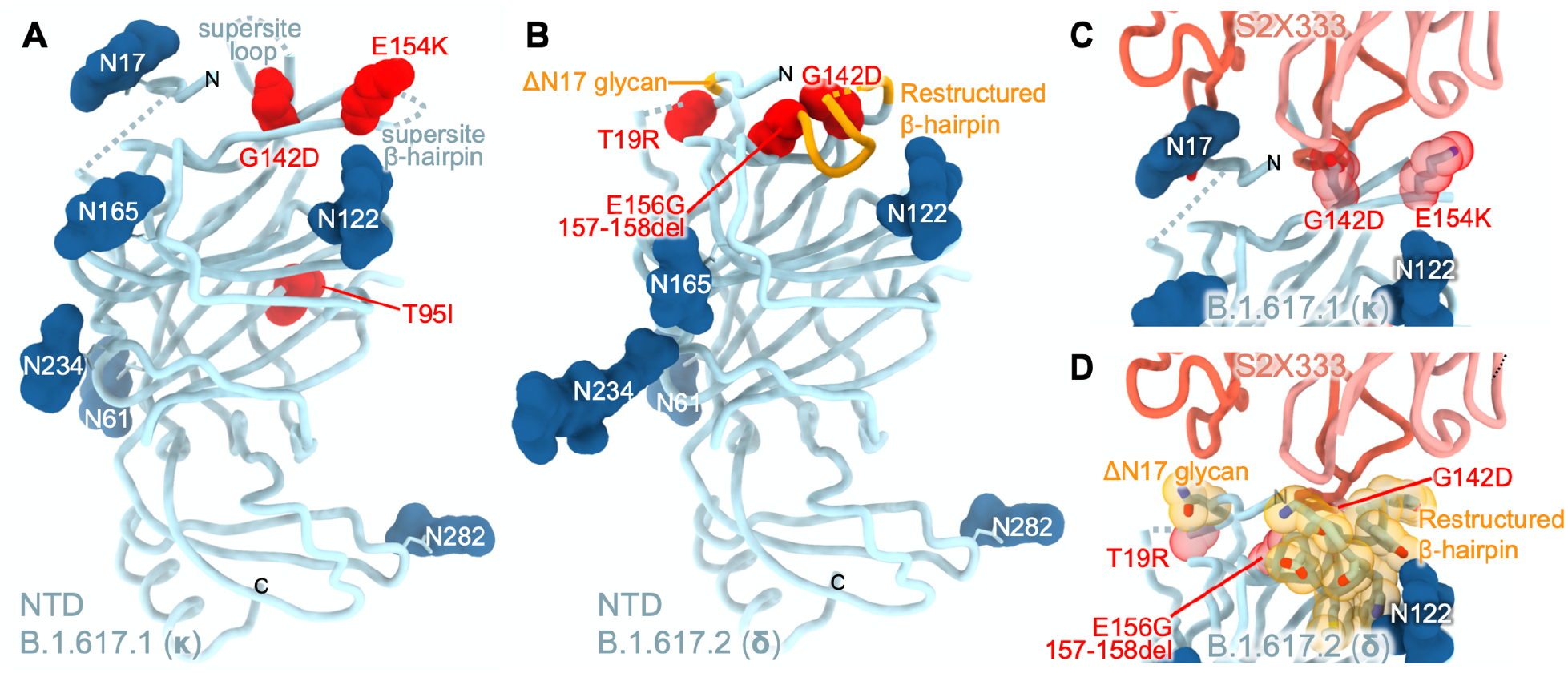
Remodeling of the NTD antigenic supersite in the B.1.617.1 and B.1.617.2 S variants. **A-B,** Ribbon diagrams of the B.1.617.1 (A) and B.1.617.2 (B) NTDs in the same orientation. Mutated residues are rendered as red spheres and N-linked glycans are shown as dark blue surfaces. Segments with notable structural changes as a consequence of these mutated residues are shown in orange and labeled. **C-D,** Zoomed-in views of the B.1.617.1 (C) and B.1.617.2 (D) NTD antigenic supersites highlighting incompatibility with recognition by the S2X333 mAb (used here as an example of prototypical NTD neutralizing mAb). N- and C-termini are labeled.

We therefore evaluated binding of a panel of neutralizing NTD antibodies to B.1.617.1, B.1.617.2, B.1.1.7, B.1.351, P.1, and B.1.427/B.1.429 S ectodomain trimers by ELISA (**Fig 4A, Fig S4**). Out of 11 neutralizing mAbs tested, we observed a 10-fold or greater reduction in binding for 8, 10, 10, 10, 3, and 11 mAbs to B.1.617.1, B.1.617.2, B.1.1.7, B.1.351, P.1, and B.1.427/B.1.429 S as a result of the introduced mutations and deletions whereas the non-neutralizing S2L20 mAb efficiently recognized all S trimers, confirming proper folding (**Fig 4A**). Collectively, these data indicate that NTD-specific neutralizing Abs exert a selective pressure participating in evolution of the antigenic supersite leading to neutralization escape in numerous variants (*62–64*), including the B.1.617.1 and B.1.617.2. The diversity of molecular solutions observed in SARS-CoV-2 variants to evade NTD-targeted neutralizing Ab responses further underscores the unparalleled plasticity of this domain.

**Figure 4.**
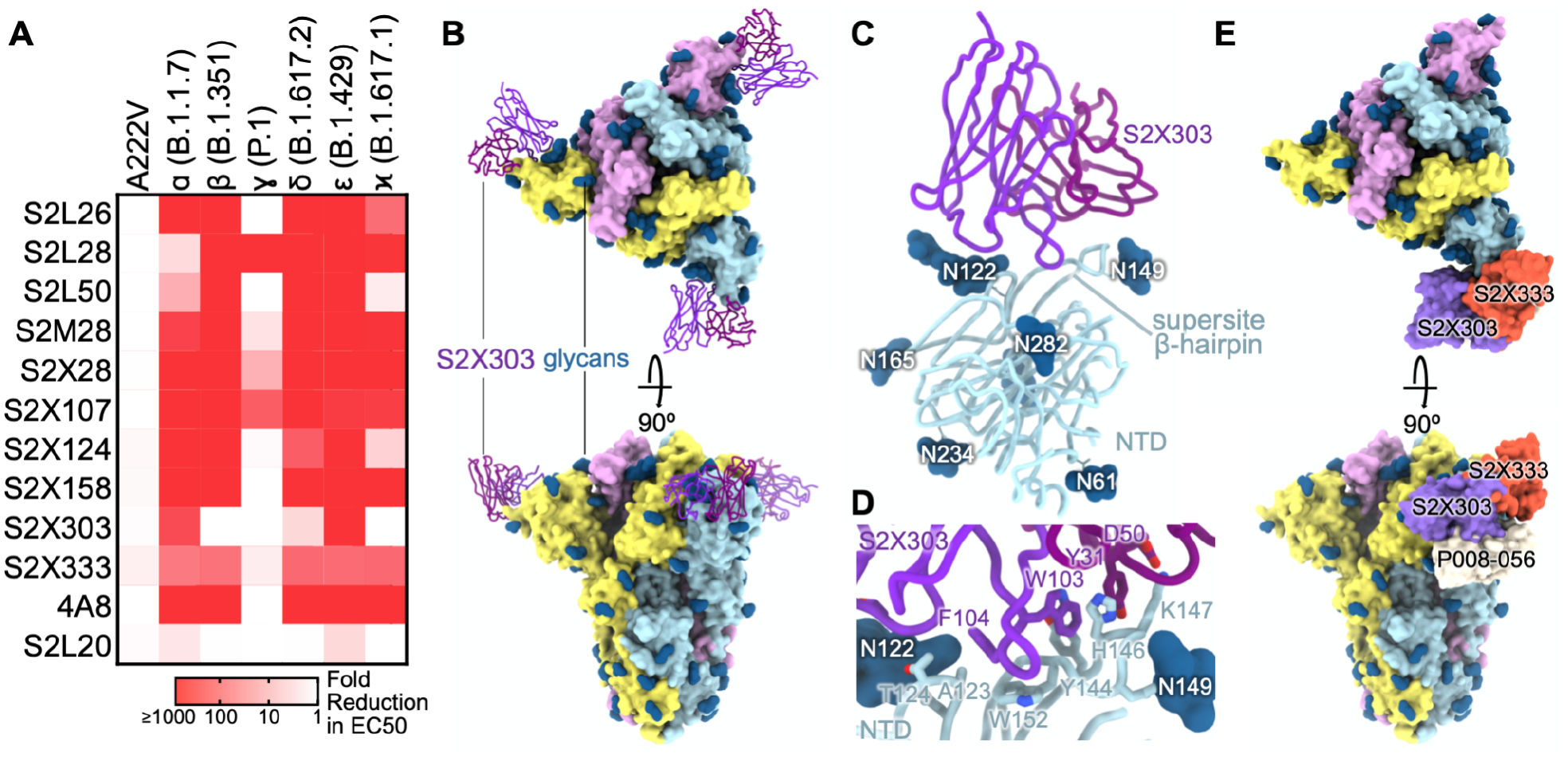
S2X303 defines a subclass of site i NTD neutralizing mAbs cross-reacting with several variants. (**A**) Binding of a panel of 11 neutralizing (antigenic site i) and 1 non-neutralizing (S2L20, antigenic site iv) NTD-specific mAbs to recombinant SARS-CoV-2 S variants analyzed by ELISA displayed as a heat map. (**B**) Structure of the B.1.617.1 S trimer (surface rendering) bound to the S2X303 Fab fragment (ribbons) shown in two orthogonal orientations. SARS-CoV-2 S protomers are colored pink, cyan, and gold, whereas the S2X303 Fab heavy and light chains are colored dark and light purple, respectively. Only the Fab variable domains are resolved and therefore modeled in the map. N-linked glycans are rendered as dark blue spheres. (**C**) Ribbon diagram of the S2X303-bound SARS-CoV-2 B.1.617.1 NTD. (**D**) Zoomed-in view of the S2X303-bound B.1.617.1 NTD with key residues involved in the interface shown as sticks. (**E**) Structure of the S trimer bound to the S2X303 overlaid with S2X333 and P008-056 antibodies (PDB IDs 7LXW and 7NTC, respectively) shown as a surface rendering. S is colored as in (B); S2X303, S2X333, and P008-056 are shown in purple, orange, and light grey, respectively.

**Fig. S4.**
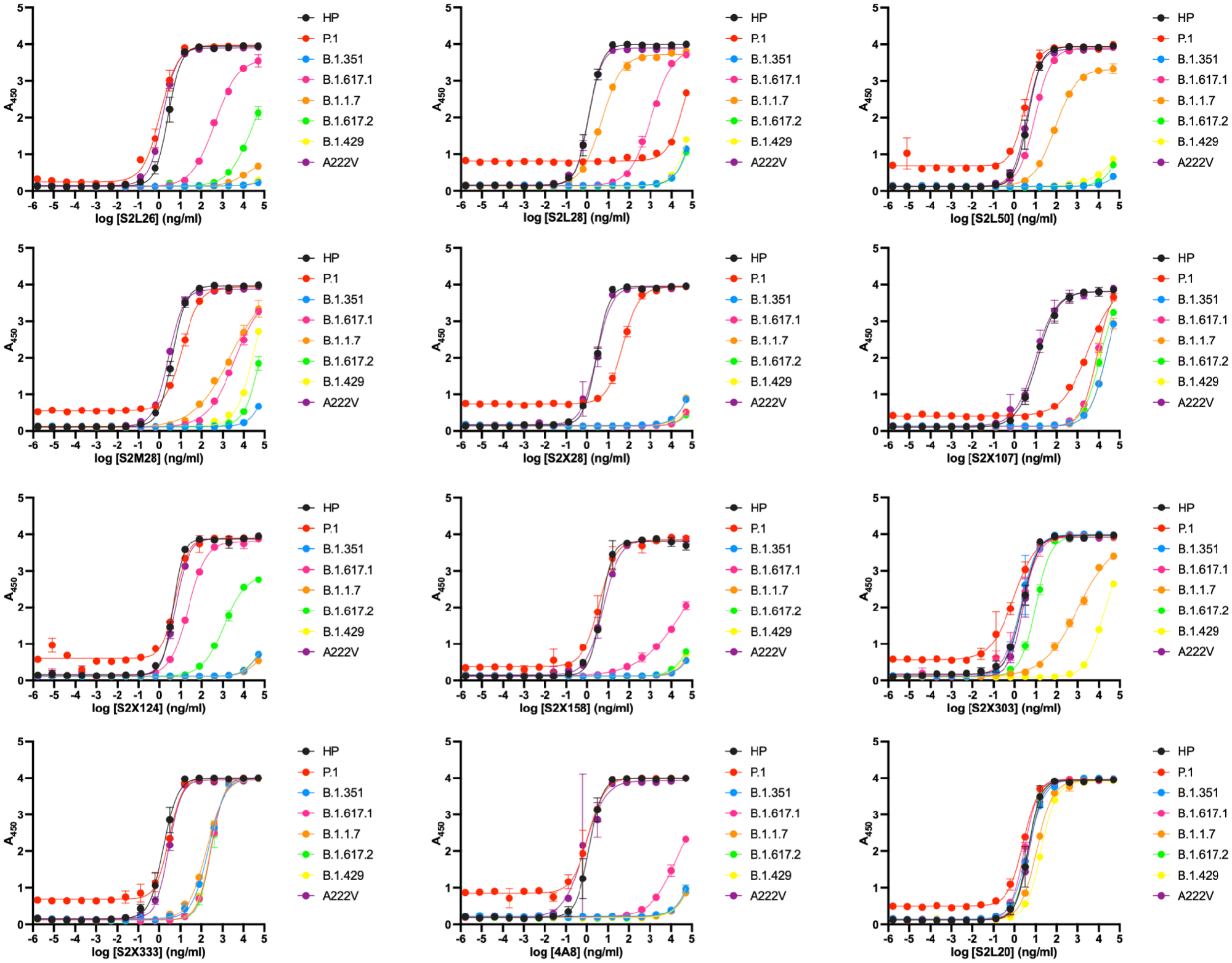
Effect of SARS-CoV-2 variant mutations on NTD-targeted mAb binding. Binding of a panel of 11 neutralizing (antigenic site i) and 1 non-neutralizing (antigenic site iv, S2L20) NTD-specific mAbs to recombinant SARS-CoV-2 variant ectodomains analyzed by ELISA.

S2X303 stood out due to its greater cross-reactivity with variants, including B.1.351, P.1, and to a lesser extent B.1.617.2, compared to all other neutralizing mAbs evaluated (**Fig 4A**). To provide a structural framework for understanding the S2X303 binding breadth, we characterized its Fab fragment bound to B.1.617.1 S using cryoEM (**Fig 4B**). Focused classification and local refinement of the S2X303-bound NTD yielded a map at 3.5 Å resolution revealing the unique recognition mode of this mAb (**Fig 4C, Fig S3G-I, Table S3**).

S2X303 recognizes the NTD with an angle of approach almost orthogonal relative to several previously described NTD-specific neutralizing mAbs and its epitope is only partially overlapping with the NTD antigenic supersite (*23*). Specifically, the S2X303 complementary determining region 3-dominated paratope exclusively contacts residues 123-125 (along with the glycan at position N122), that are part of the NTD galectin-like distal β-sheet, and residues 144-154 within the supersite β-hairpin (**Fig 4D-E**). As a result, S2X303 defines a new class of NTD-specific mAbs that differs from S2X333 (*23*), a canonical ultrapotent NTD-specific mAb, and from P008_056 (*65*), which interferes with biliverdin binding, with respect to their footprints and angles of approach (**Fig 4E**). Moreover, S2X303 exhibits unprecedented cross-reactivity compared to all other mAbs and can bind to a large number of variants (**Fig 4A**).

Mutations found in the B.1.617.1 and B.1.617.2 variants mediate immune evasion by eroding infection- and vaccine-elicited serum neutralizing Ab titers due to structural alteration present in major antigenic sites within the RBD and NTD (*37, 66, 67*). Neither of the fusion machinery B.1.617.1 Q1071H or the B.1.617.2 D950N substitutions are part of epitopes known to be recognized by neutralizing Abs. The B.1.617.2+ variant is associated with a severe dampening of neutralizing Ab titers due to the additional K417N RBD substitution, also found in the B.1.351 variant of concern (*1, 2, 38*). However, the deleterious effect of this mutation on ACE2 binding and absence of the compensatory N501Y mutation found in B.1.351 (*4*) might be associated with a fitness cost, putatively explaining the small number of genomes detected for this variant. Based on the roughly comparable ACE2 binding affinities of the B.1.617.1, B.1.617.2 and the ancestral RBDs, we propose that other factors contribute to the enhanced transmissibility of the B.1.617.2 variant. The P681R mutation found in the B.1.617.1 and B.1.617.2 variants was recently shown to enhance cleavage at the S_1_/S_2_ boundary and cell-cell fusion (*37, 67, 68*). Since the S_1_/S_2_ cleavage site is key for transmission and pathogenicity (*69, 70*), it appears likely that this mutation contributes to the success of these lineages. Furthermore, the L452R mutation, which augments stability/expression (*59*), increases viral replication kinetics relative to the ancestral virus (*71*). The enhanced ability of B.1.1.7 to antagonize host innate immunity through upregulation of orf6 and orf9b was suggested to have participated in the success of this variant (*72*). It is possible that evolved a similar strategy to reach global domination warranting further studies to uncover the contribution of innate immune antagonism to the continued emergence of variants with greater transmissibility.

We demonstrate here that S309, the parent mAb of sotrovimab which has received an emergency use authorization from the FDA, is unaffected by antigenic drift observed in variants of concern and interest due to recognition of a conserved RBD epitope (*12, 15, 56*). The recent discovery of multiple additional conserved antigenic sites recognized by RBD-specific mAbs with (near) pan-sarbecovirus neutralizing activity (*16, 17, 21, 73–75*) and the anticipated continued emergence of SARS-CoV-2 variants motivate the clinical development and deployment of some of these mAbs for prophylaxis with at-risk patients and for treatments of unvaccinated individuals or breakthrough infections. Moreover, next-generation vaccine candidates have recently been described to elicit broad sarbecovirus immunity (*41, 76, 77*), holding the promise to be resilient to the emergence of SARS-CoV-2 variants and of new zoonotic sarbecoviruses. Looking forward, the discovery of broadly neutralizing mAbs targeting the fusion machinery makes tangible the development of a universal β-coronavirus vaccine (*27–30, 78*).

---

**Table S1:**
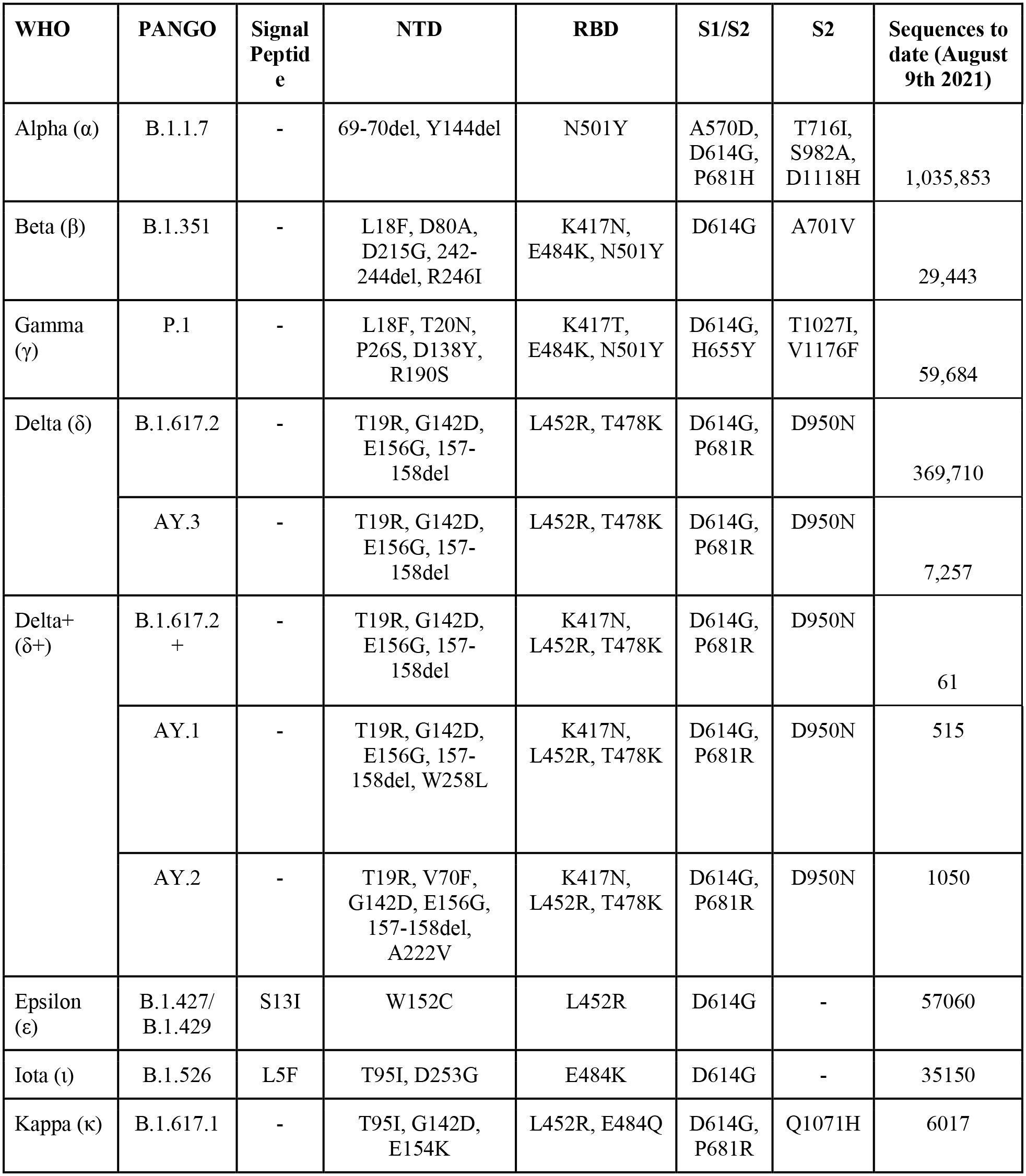
Summary of mutations present in select SARS-CoV-2 S variants.

| WHO | PANGO | Signal Peptide | NTD | RBD | S1/S2 | S2 | Sequences to date (August 9th 2021) |
| --- | --- | --- | --- | --- | --- | --- | --- |
| Alpha ($\alpha$) | B.1.1.7 | - | 69-70del, Y144del | N501Y | A570D, D614G, P681H | T716I, S982A, D1118H | 1,035,853 |
| Beta ($\beta$) | B.1.351 | - | L18F, D80A, D215G, 242-244del, R246I | K417N, E484K, N501Y | D614G | A701V | 29,443 |
| Gamma ($\gamma$) | P.1 | - | L18F, T20N, P26S, D138Y, R190S | K417T, E484K, N501Y | D614G, H655Y | T1027I, V1176F | 59,684 |
| Delta ($\delta$) | B.1.617.2 | - | T19R, G142D, E156G, 157-158del | L452R, T478K | D614G, P681R | D950N | 369,710 |
|  | AY.3 | - | T19R, G142D, E156G, 157-158del | L452R, T478K | D614G, P681R | D950N | 7,257 |
| Delta+ ($\delta$+) | B.1.617.2 + | - | T19R, G142D, E156G, 157-158del | K417N, L452R, T478K | D614G, P681R | D950N | 61 |
|  | AY.1 | - | T19R, G142D, E156G, 157-158del, W258L | K417N, L452R, T478K | D614G, P681R | D950N | 515 |
|  | AY.2 | - | T19R, V70F, G142D, E156G, 157-158del, A222V | K417N, L452R, T478K | D614G, P681R | D950N | 1050 |
| Epsilon ($\epsilon$) | B.1.427/ B.1.429 | S13I | W152C | L452R | D614G | - | 57060 |
| Iota ($\iota$) | B.1.526 | L5F | T95I, D253G | E484K | D614G | - | 35150 |
| Kappa ($\kappa$) | B.1.617.1 | - | T95I, G142D, E154K | L452R, E484Q | D614G, P681R | Q1071H | 6017 |

**Table S2:**
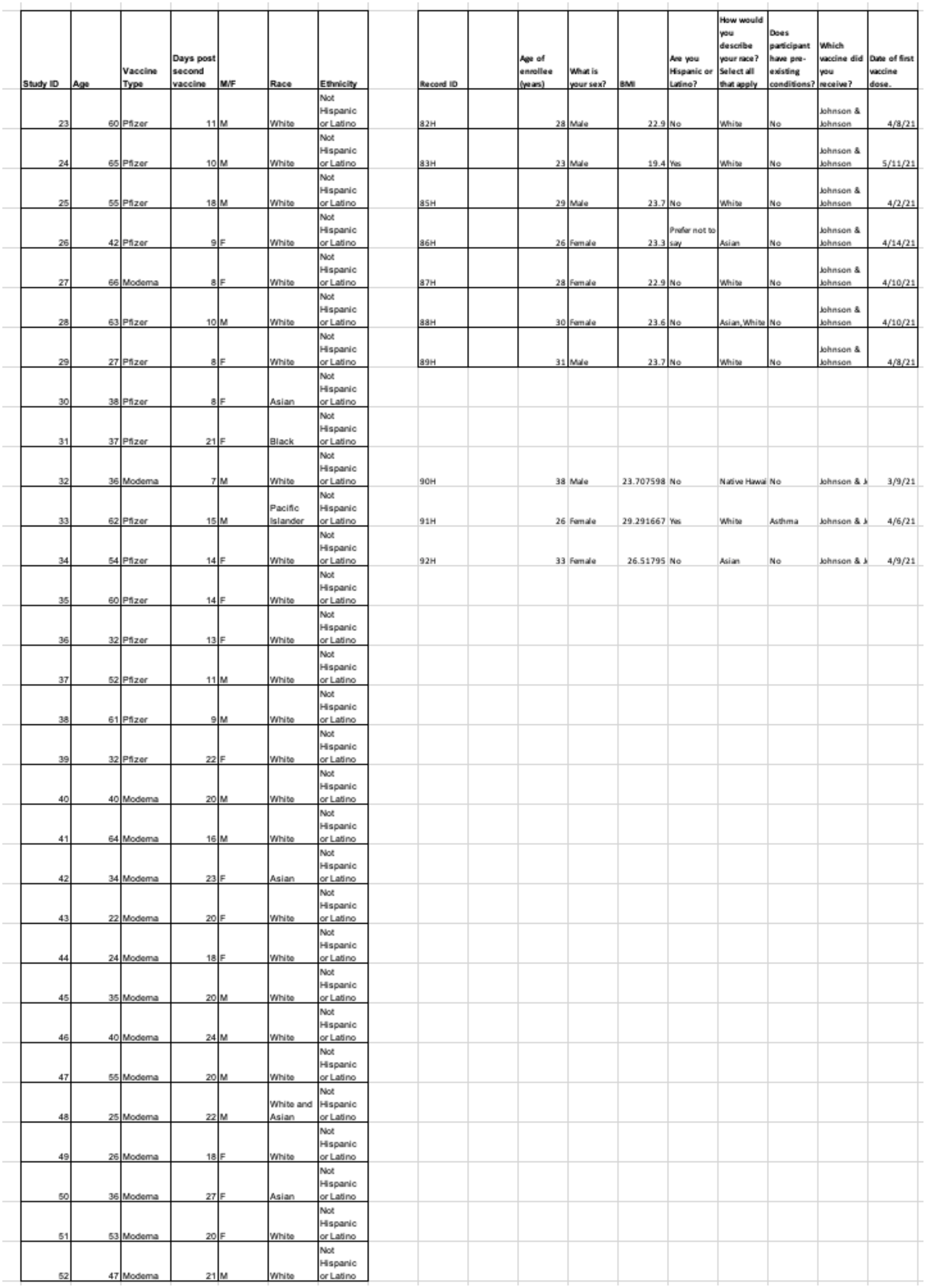
Demographics of vaccinated individuals.

**Table S3.**
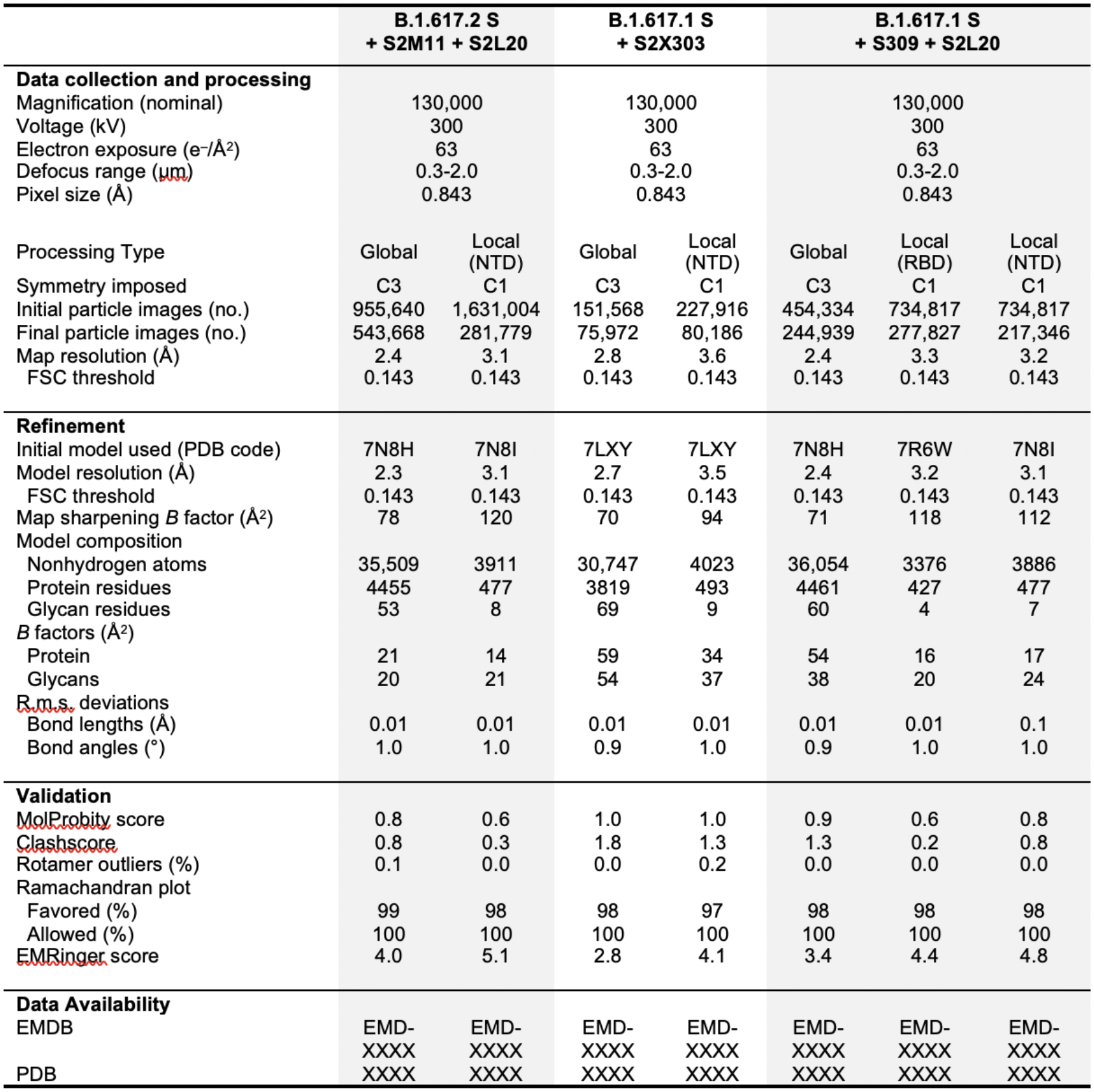
Cryo-EM data collection, refinement and validation statistics.

**Table S4:** Binding kinetics of RBD to ACE2.

| **ELISA** | **EC50 (nM) ± SEM** |
| --- | --- |
| **WT** | $5 \pm 1$ |
| **B.1.1.7** | $0.48 \pm 0.02$ |
| **B.1.617.1** | $9 \pm 1$ |
| **B.1.617.2** | $5 \pm 2$ |
| **B.1.617.2+** | $40 \pm 20$ |

| **SPR** | **K<sub>D</sub> (nM) ± SEM** | **k<sub>on</sub> (M<sup>-1</sup>s<sup>-1</sup>)** | **k<sub>off</sub> (s<sup>-1</sup>)** |
| --- | --- | --- | --- |
| **WT** | $78 \pm 8$ | $7.7 \times 10^4$ | $6.7 \times 10^{-3}$ |
| **B.1.1.7** | $15.0 \pm 0.4$ | $7.5 \times 10^4$ | $1.2 \times 10^{-3}$ |
| **B.1.617.1** | $71 \pm 3$ | $6.0 \times 10^4$ | $4.8 \times 10^{-3}$ |
| **B.1.617.2** | $63 \pm 3$ | $5.9 \times 10^4$ | $4.3 \times 10^{-3}$ |
| **B.1.617.2+** | $183 \pm 4$ | $6.6 \times 10^4$ | $1.3 \times 10^{-2}$ |

| **BLI** | **K<sub>D</sub> (nM) ± SEM** | **k<sub>on</sub> (M<sup>-1</sup>s<sup>-1</sup>)** | **k<sub>off</sub> (s<sup>-1</sup>)** |
| --- | --- | --- | --- |
| **WT** | $147 \pm 3$ | $9.5 \times 10^4$ | $1.4 \times 10^{-2}$ |
| **B.1.1.7** | $26 \pm 4$ | $1.3 \times 10^5$ | $2.8 \times 10^{-3}$ |
| **B.1.617.1** | $88 \pm 1$ | $1.0 \times 10^5$ | $9.1 \times 10^{-3}$ |
| **B.1.617.2** | $180 \pm 30$ | $7.4 \times 10^4$ | $1.1 \times 10^{-2}$ |
| **B.1.617.2+** | $550 \pm 80$ | $5.1 \times 10^4$ | $2.4 \times 10^{-2}$ |

## Author contributions

M.M. and D.V conceived the project. M.M., A.C.W. and D.V designed experiments. M.M analyzed the incidence of variants. M.M. and J.E.B. expressed and purified proteins and performed ELISAs. M.M. carried out cryoEM sample preparation, data collection, processing and model building and refinement. A.C.W carried out BLI analysis. L.R. performed SPR analysis. A.C.W. and K.S. performed pseudovirus neutralization assays and analysis. H.V.D., A.M., S.W.T., M.Mi., L.C., M.S.P., H.Y.C, W.C.V.V, and D.C. provided unique reagents, G.S, D.C. and D.V. supervised experiments. M.M. and D.V wrote the manuscript with input from all authors.

## Acknowledgements

We thank Hideki Tani (University of Toyama) for providing the reagents necessary for preparing VSV pseudotyped viruses. This study was supported by the National Institute of Allergy and Infectious Diseases (DP1AI158186 and HHSN272201700059C to D.V.), a Pew Biomedical Scholars Award (D.V.), an Investigators in the Pathogenesis of Infectious Disease Awards from the Burroughs Wellcome Fund (D.V.), Fast Grants (D.V.) and the Bill & Melinda Gates Foundation (OPP1156262 to D.V.), the University of Washington Arnold and Mabel Beckman cryoEM center and the National Institute of Health grant S10OD032290 (to D.V.) and grant U01 AI151698 for the United World Antiviral Research Network (UWARN) as part of the Centers for Research in Emerging Infectious Diseases (CREID) Network. We thank Julia L. Mullen, Ginger Tsueng, Alaa Abdel Latif, Manar Alkuzweny, Marco Cano, Emily Haag, Jerry Zhou, Mark Zeller, Emory Hufbauer, Nate Matteson, Kristian G. Andersen, Chunlei Wu, Andrew I. Su, Karthik Gangavarapu, Laura D. Hughes, and the Center for Viral Systems Biology *outbreak.info*. (https://outbreak.info/).

## Methods

### Variant incidence analysis

Average daily prevalence for B.1.1.7, B.1.351, P.1, B.1.617.2 (including AY.3), B.1.526, B.1.427/B.1.429, and B.1.617.2+ (including AY.1 and AY.2) were obtained from GISAID (using outbreak.info) and plotted using GraphPad PRISM software (version 9.2.0).

### Sample donors

Blood samples were collected from participants who had received both doses of the Pfizer’s BNT162b2 vaccine or Moderna’s mRNA-1273 vaccine and were 7–30 days post second dose. Individuals were enrolled in the UWARN: COVID-19 in WA study at the University of Washington in Seattle, WA. This study was approved by the University of Washington Human Subjects Division Institutional Review Board (STUDY00010350). The J&J/Janssen Ad26.COV2.S vaccine samples were collected as part of the HAARVI study and was approved by the University of Washington Human Subjects Division Institutional Review Board (STUDY00000959). Baseline sociodemographic and clinical data for these individuals are summarized in Table S2.

### Recombinant expression of mAbs

Recombinant mAbs were expressed in ExpiCHO cells at 37 °C and 8% CO2. Cells were transfected using ExpiFectamine. Transfected cells were supplemented 1 day after transfection with ExpiCHO Feed and ExpiFectamine CHO Enhancer. Cell culture supernatant was collected eight days after transfection and filtered through a 0.2 μm filter. Recombinant antibodies were affinity purified on an ÄKTA xpress FPLC device using 5 mL HiTrapTM MabSelectTM PrismA columns followed by buffer exchange to Histidine buffer (20 mM Histidine, 8% sucrose, pH 6) using HiPrep 26/10 desalting columns.

### Variant construct generation

The WT, B.1.1.7, B.1.617.1, B.1.617.2, and B.1.617.2+ SARS-CoV-2 RBD construct were synthesized by GenScript into pCMVR with an N-terminal mu-phosphatase signal peptide, and a C-terminal octa-histidine tag (GHHHHHHHH) and an avi tag (GLNDIFEAQKIEWHE)). The boundaries of the construct are N-328-RFPN-331 and 528-KKST-531.

The hACE2 construct was previously synthesized by Twist into pTwist+CMV(residues 1-615 with a C-terminal avi tag-10xHis-GGG-tag, and N-terminal signal peptide) (*59*).

The SARS-CoV-2 S ectodomain with hexapro mutations (*45*), S383C/D985C mutations (*46*), the native furin cleavage site (RRAR), and B.1.617.1 spike mutations (T95I, G142D, E154K, L452R, E484Q, D614G, P681R and Q1071H) was synthesised by GenScript into pCMV with a C-terminal foldon and avi tag followed by an octa-histidine tag. The SARS-CoV-2 S ectodomain with VFLIP mutations (*48*), the native furin cleavage site (RRAR), and B.1.617.2 spike mutations (T19R, G142D, E156G, T478K, and D950N substitutions and a deletion of residues 157 and 158) was synthesised by GenScript into pCMV with a C-terminal avi tag followed by an octa-histidine tag. The SARS-CoV-2 S ectodomain with hexapro mutations and P.1 spike mutations (L18F, T20N, P26S, D138Y, R190S, K417T, E484K, N501Y, D614G, H655Y, T1027I, and V1176F) was synthesised by GenScript into pCMVR with foldon, a C-terminal avi tag followed by an octa-histidine tag. The SARS-CoV-2 S ectodomain with the 2P mutations and B.1.1.7 spike mutations (del69/70, del144/145, N501Y, A570D, D614G, P681H, T716I, S982A, and D1118H) was synthesised by GenScript into pCMV with a C-terminal foldon and avi tag followed by an octa-histidine tag. The SARS-CoV-2 S ectodomain with the 2P mutations and B.1.351 spike mutations (D80A, D215G, 242-244del, R246I, K417N, E484K, N501Y, D614G, and A701V) was synthesised by GenScript into pCMV with a C-terminal foldon and avi tag followed by an octa-histidine tag.

### Production of recombinant glycoproteins

hACE2 and the biotinylated RBDs were produced in 25 mL cultures of Expi293F Cells (ThermoFisher Scientific) grown in suspension using Expi293 Expression Medium (ThermoFisher Scientific) at 37°C in a humidified 5% CO_2_ incubator rotating at 130 rpm. Cells grown to a density of 3 million cells per mL were transfected using the ExpiFectamine 293 Transfection Kit (ThermoFisher Scientific) and cultivated for four days. Proteins were purified from clarified supernatants using a nickel HisTrap HP affinity column (Cytiva) and washed with ten column volumes of 20 mM imidazole, 25 mM sodium phosphate pH 8.0, and 300 mM NaCl before elution with a gradient of 500 mM imidazole. Proteins were buffer exchanged into 20 mM sodium phosphate pH 8 and 100 mM NaCl and concentrated using 30 kDa or 10 kDa centrifugal filters (Amicon Ultra, MilliporeSigma) for hACE2 and the biotinylated RBDs, respectively, before being flash frozen.

The SARS-CoV-2 S ectodomains were produced in 100 mL cultures of Expi293F Cells (ThermoFisher Scientific) grown in suspension using Expi293 Expression Medium (ThermoFisher Scientific) at 37°C in a humidified 8% CO_2_ incubator rotating at 130 rpm. Cells grown to a density of 2.5 million cells per mL were transfected using the ExpiFectamine 293 Transfection Kit (ThermoFisher Scientific) and cultivated for four days at which point the supernatant was harvested. S ectodomains were purified from clarified supernatants using a Cobalt affinity column (Cytiva, HiTrap TALON crude), washing with 20 column volumes of 20 mM Tris-HCl pH 8.0 and 150 mM NaCl and eluted with a gradient of 600 mM imidazole. The S ectodomain was then concentrated using a 100 kDa centrifugal filter (Amicon Ultra 0.5 mL centrifugal filters, MilliporeSigma), residual imidazole was washed away by consecutive dilutions in the centrifugal filter unit with 20 mM Tris-HCl pH 8.0 and 150 mM NaCl, and finally concentrated to 2 mg/mL and flash frozen.

### Production of VSV pseudovirus

SARS-CoV-2 D614G, B.1.617.1 (mutations T95I, G142D, E154K, L452R, E484Q, D614G, P681R and Q1071H), B.1.617.2 (mutations T19, G142D, E156G, 157-158del, T478K, D950N) and B.1.617.2+ (B.1.617.2 mutations plus K417N) pseudotypes were prepared similarly as previously described (*47*). Briefly, HEK-293T cells seeded in poly-D-lysine coated 100 mm dishes at ∼75 % confluency were washed five times with Opti-MEM and co-transfected with Lipofectamine 2000 (Life Technologies) with 24 µg of the respective S glycoprotein plasmids. After 5 h at 37°C, media supplemented with 20% FBS and 2% PenStrep was added. After 20 hours, cells were washed five times with DMEM and cells were transduced with VSVΔG-luc (*39*) and incubated at 37°C. After 2 h, infected cells were washed an additional five times with DMEM prior to adding media supplemented with anti-VSV-G antibody (I1-mouse hybridoma supernatant diluted 1:25, from CRL-2700, ATCC) to reduce parental background. After 18-24 h, the supernatant was harvested and clarified by low-speed centrifugation at 2,500 g for 10 min. The supernatant was then filtered (0.45 μm) and concentrated 10 times using a 30 kDa cut off membrane. The pseudotypes were then aliquoted and frozen at −80 °C.

### VSV neutralization

To evaluate neutralization of D614G, B.1.617.1, B.1.617.2 and B.1.617.2 + pseudotypes by sera of individuals immunized with Pfizer/BioNtech BNT162b2, Moderna mRNA1273 and Janssen Ad26.COV2.S, HEK-293T cells expressing hACE2 (*40*) in DMEM supplemented with 10% FBS and 1% PenStrep were seeded at 20,000 cells per well into clear bottom, white manually poly-D-lysine coated 96 well plates and incubated at 37°C.

The following day, an additional half-area, 96-well plate was prepared with eight 3-fold serial dilutions of sera from Pfizer/BioNtech BNT162b2, Moderna mRNA1273 or Janssen Ad26.COV2.S vaccine recipients. An equal volume of each 1:25 DMEM diluted D614G, B.1.617.1, B.1.617.2 and B.1.617.2+ pseudovirus were then added to the half-area plate. The mixture was then incubated at room temperature for 30 minutes.

At this stage, excess DMEM was removed from the cells and 40 µL from each well (containing sera and pseudovirus) was transferred to the 96-well plate seeded with HEK-293T cells expressing hACE2 and incubated at 37°C for 2 h. After 2 h, an additional 40 µL of DMEM supplemented with 20% FBS and 2% PenStrep was added to the cells.

After 20 h, 40 µL of One-Glo-EX substrate (Promega) was added to each well and incubated on a plate shaker in the dark for 5 min. The plates were immediately read on a Biotek plate reader. Measurements were done in duplicate with biological replicates. Relative luciferase units were plotted and normalized in Prism (GraphPad): cells alone without pseudovirus was defined as 0 % infection, and cells with virus only (no sera) was defined as 100 % infection. Prism (GraphPad) nonlinear regression with “[inhibitor] versus normalized response with a variable slope” was used to determine IC50 values from curve fits. Means of these duplicates were compared against G614 by two-way ANOVA (Dunnet’s test) using GraphPad PRISM software (version 9.2.0).

### Western Blot

Undiluted pseudoviruses were added to 4X SDS-PAGE loading buffer. Samples were run on a 4%–15% gradient Tris-Glycine Gel (BioRad) and transferred to PVDF membranes. An anti-S_2_ SARS-CoV-2 S polyclonal primary antibody (1:1,500 dilution,Invitrogen PA5-114534) and an Alexa Fluor 680-conjugated goat anti-rabbit secondary antibody (1:20,000 dilution, Jackson Laboratory 111-625-144) were used for Western-blotting. A LI-COR processor was used to develop images.

### ELISA

For ELISA experiments with NTD-targeted mAbs, 384-well Maxisorp plates (ThermoFisher Scientific 464718) were coated overnight at 4°C with 2 μg/mL of S glycoprotein in 20mM HEPES pH 8 and 150mM NaCl. Plates were slapped dry and blocked with Blocker Casein in TBS (ThermoFisher Scientific 37532) for one hour at 37°C. Plates were slapped dry and mAbs were serially diluted 1:5 in TBST with an initial concentration of 50 µg/ml. Plates were left for one hour at 37°C and washed 4x with TBST, then 1:5000 Goat anti-Human (ThermoFisher Scientific A18817) was added. Plates were left for one hour at 37°C and washed 4x with TBST, and then TMB Microwell Peroxidase (Seracare 5120-0083) was added. The reaction was quenched after 4 minutes with 1 N HCl and the A450 of each well was read using a BioTek plate reader.

For ELISA experiments with RBDs, 384-well Maxisorp plates (ThermoFisher Scientific 464718) were coated overnight at 4°C with 4 µg/mL of hACE2-His in 20mM Sodium Phosphate pH 8 and 100mM NaCl. Plates were slapped dry and blocked with Blocker Casein in TBS (ThermoFisher Scientific 37532) for one hour at 37°C. Plates were slapped dry and wild-type RBD, B.1.1.7 RBD, B.1.351 RBD, B.1.617.1 RBD, B.1.617.2 RBD, and B.1.617.2+ RBD were serially diluted 1:3 in TBST with an initial concentration of 1542nM. Plates were left for one hour at 37°C, then washed 4x with TBST using a 405 TS Microplate Washer (BioTek) followed by addition of 2 μg/mL S309 mAb (*15*). Plates were left for one hour at 37°C and washed 4x with TBST, then 1:5000 Goat anti-Human (ThermoFisher Scientific A18817) was added. TMB Microwell Peroxidase (Seracare 5120-0083) was added after another hour at 37°C and 4x wash with TBST. The reaction was quenched after 1-2 minutes with 1 N HCl and the A450 of each well was read using a BioTek plate reader.

### CryoEM sample preparation and data collection

50 μL of 2 mg/mL SARS-CoV-2 S B.1.617.1 ectodomain was incubated with 29 µl 3.4 mg/ml S309 Fab and 2.2 μL of 67 mg/mL S2L20 Fab in 150 mM NaCl and 20 mM Tris-HCl pH 8 for 30 min at 37°C. Alternatively, 50 μL of 2 mg/mL SARS-CoV-2 S B.1.617.2 ectodomain was incubated with 34 µl 2.9 mg/ml S2M11 Fab in 150 mM NaCl and 20 mM Tris-HCl pH 8 for 10 min at 37°C, and then 2.2 μL of 67 mg/mL S2L20 Fab was added for 20 min at 37°C. Alternatively, 50 μL of 2 mg/mL SARS-CoV-2 S B.1.617.1 ectodomain was incubated with 3.6 µl 28 mg/ml S2X303 Fab in 150 mM NaCl and 20 mM Tris-HCl pH 8 for 30 min at 37°C.

Unbound Fab was then washed away with six consecutive dilutions in 400 μL of 20 mM Tris-HCl pH 8.0 and 150 mM NaCl over a 100 kDa centrifugal filter (Amicon Ultra 0.5 mL centrifugal filters, MilliporeSigma). The complex was concentrated to 3.6 mg/mL and 3 μL was immediately applied onto a freshly glow discharged 2.0/2.0 UltraFoil grid (200 mesh), plunge frozen using a vitrobot MarkIV (ThermoFisher Scientific) using a blot force of −1 and 6.5 s blot time at 100% humidity and 23°C.

Data were acquired using the Leginon software (*79*) to control a FEI Titan Krios transmission electron microscope equipped with a Gatan K3 direct detector and operated at 300 kV with a Gatan Quantum GIF energy filter. The dose rate was adjusted to 3.75 counts/super-resolution pixel/s, and each movie was acquired in 75 frames of 40 ms with a pixel size of 0.843 Å and a defocus range comprised between −0.2 and −2.0 μm.

### CryoEM data processing

Movie frame alignment, estimation of the microscope contrast-transfer function parameters, particle picking and extraction (with a downsampled pixel size of 1.686 Å and box size of 256 pixels^2^) were carried out using Warp (*80*). Reference-free 2D classification was performed using cryoSPARC (*81*) to select well-defined particle images. 3D classification with 50 iterations each (angular sampling 7.5 ° for 25 iterations and 1.8 ° with local search for 25 iterations) were carried out using Relion without imposing symmetry. 3D refinements were carried out using non-uniform refinement in cryoSPARC (*82*) before particle images were subjected to Bayesian polishing using Relion (*83*) during which particles were re-extracted with a box size of 512 Å at a pixel size of 0.843 Å. Next, 86 optics groups were defined based on the beamtilt angle used for data collection. Another round of non-uniform refinement in cryoSPARC was then performed concurrently with global and per-particle defocus refinement. For focused classification, particles were symmetry-expanded in Relion (*84, 85*), and the particles were 3D classified in Relion without alignment using a mask that encompasses part of the NTD and the S2L20 VH/VL region, or the RBD and the S309 VH/VL region. Particles in well-formed 3D classes were then used for local refinement in cryoSPARC. Reported resolutions are based on the gold-standard Fourier shell correlation of 0.143 criterion and Fourier shell correlation curves were corrected for the effects of soft masking by high-resolution noise substitution (*86, 87*).

### CryoEM model building and analysis

UCSF Chimera (*88*) and Coot (*89*) were used to fit atomic models of S2M11, S2L20, S309, and SARS-CoV-2 S (PDB 7LXY, 7N8I, 7R6W) into the cryo-EM maps. The model was then refined and rebuilt into the map using Coot (*89*), Rosetta (*90, 91*), Phenix (*92*), and ISOLDE (*93*). Model validation and analysis used MolProbity (*94*), EMringer (*95*), Phenix (*92*) and Privateer (*96*). Figures were generated using UCSF ChimeraX (*97*).

### ACE2 binding measurements using Biolayer interferometry

His-avi-tagged wildtype, B.1.1.7 (N501Y) B.1.617.1 (L452R, E484Q), B.1.617.2 (L452R, T478K), or B.1.617.2+ (K417N, L452R, T478K) RBD were biotinylated and immobilized at 5 ng/μL in undiluted 10X kinetics buffer (Pall) to SA sensors that were pre-hydrated in water for at least 10 minutes and then equilibrated into 10X Kinetics Buffer (Pall). The RBDs were loaded to a level of 1nm total shift. The loaded tips were then dipped into a dilution series of monomeric ACE2-his in 10X Kinetics Buffer (Pall) starting at 1000 or 5000 nM for 300 seconds prior to 300 seconds dissociation in 10X Kinetics buffer for kinetics determination. The data were baseline subtracted and the plots fitted using the Pall FortéBio/Sartorius analysis software (v.12.0). Data were plotted in Graphpad Prism (v.9.0.2). These experiments were done side-by-side with two separate RBD protein preparations and two separate ACE2 preparations and a representative experiment is shown.

### ACE2 binding measurements using surface plasmon resonance

Measurements were performed using a Biacore T200 instrument. The Cytiva Biotin CAPture Kit, Series S, was used for surface capture of biotinylated RBDs. Running buffer was HBS-EP+ pH 7.4 (Cytiva) and measurements were performed at 25 °C. Experiments were performed with a 3-fold dilution series of monomeric hACE2: 200 nM, 67 nM, 22 nM, 7.4 nM. Association was 300 s and dissociation was 450 s. Data were double reference-subtracted and fit to a 1:1 binding model using Biacore Evaluation software.

